# Data fusion for abundance estimation: community science augments systematically collected removal-in-time distance sampling data

**DOI:** 10.1101/2021.05.02.442379

**Authors:** Maxwell B. Joseph, David C. Pavlacky, Anne M. Bartuszevige

## Abstract

Ecologists use a variety of systematically and opportunistically sampled count data to estimate bird abundance, and integrating or fusing different datasets has emerged as a critical challenge in recent years. While previous work provides data integration methodology for occupancy (presence/absence) estimation, methods for abundance estimation that account for imperfect detection and disparate survey protocols remains an active area of research. Here we show how to integrate systematically collected removal-in-time distance sampling data from the Integrated Monitoring in Bird Conservation Regions (IMBCR) program with North American Breeding Bird Survey (BBS) point counts and eBird community science observations. Using the Grasshopper Sparrow (*Ammodramus savannarum*) in the Great Plains of the United States as a focal species, we demonstrate that BBS and eBird data improve predictive performance for IMBCR count data, providing more spatially refined and precise estimates of abundance at regional scales. Data fusion increased predictive performance even despite relatively weak spatial correlations among data sets. The methodology developed here provides a principled way to fuse data when estimating abundance with distance sampling, that accounts for imperfect detection and variable effort.

## 1 Introduction

A wealth of systematically and opportunistically collected data provide information about bird abundance at continental scales. Integrating these diverse datasets is difficult because of differences in survey protocols, sampling designs, and data quality. These challenges motivate recent data fusion methods, which integrate different datasets to fill spatial coverage gaps and improve species distribution models (Hudson et al. 2017). In particular, data fusion provides a framework for borrowing strengths from different data sources to improve inference about ecological processes (Hanks, Hooten, and Baker 2011). At one end of the spectrum, structured data collection designs provide representative sampling and protocols for addressing imperfect detection, and at the other end of the spectrum, less intensive inventory surveys provide additional information on spatial heterogeneity and extent. For example, Pacifici et al. (2017) combined observations from the North American Breeding Bird Survey (BBS) and eBird, and demonstrated that models using both datasets produced more accurate and precise occupancy estimates. In some applications density may be of greater interest than occupancy, and developing methods to integrate density data is an active area of research Isaac et al. (2020). Here, we fuse disparate data sources for density estimation, augmenting removal-in-time distance sampling data from the Integrated Monitoring in Bird Conservation Regions (IMBCR, Pavlacky Jr et al. (2017)) program with BBS point counts and eBird checklists.

Bird data sources differ in quality and quantity. The IMBCR program conducts bird point counts during the breeding season using a point-transect distance sampling with removal-in-time protocol. This protocol along with a stratified random sampling design provides spatially balanced information on density, availability, and detectability (Pavlacky Jr et al. 2017). IMBCR data are high quality, with representative sampling and protocols for estimating incomplete detection, but relatively low in quantity and spatial coverage (each survey costs ≈ $1200 USD). In contrast, large quantities of lower or unknown quality crowdsourced eBird data are available with potential detection bias and non-representative sampling, especially near densely populated areas (Sullivan et al. 2014). Finally, BBS data provide moderate or intermediate quality bird observations with potential detection bias and non-representative sampling along roadsides, with a decades long historical record and intermediate spatial density across much of North America (John R. Sauer et al. 2017).

Here, we test whether eBird and BBS data can be used to augment high-quality IMBCR distance sampling data that provide information on bird density and detection. We build models that (1) capitalize on the availability, detection, and density information contained in the IMBCR survey protocol, (2) borrow spatial information from BBS and eBird by combining methodological elements of distance sampling to estimate density and detectability (Amundson, Royle, and Handel 2014), and correlated spatial fields for data integration (Pacifici et al. 2017), and (3) estimate habitat relationships that may be useful for predicting species distributions. We hypothesized that the combination of IMBCR, BBS, and eBird data would provide the best performance. Specifically, we predict that including information from eBird and BBS with a structured model for IMBCR data provides the best predictive performance on withheld IMBCR data, resulting in more spatially refined and precise predictions of density from the IMBCR program.

## 2 Methods

### 2.1 Study region and focal species

The shortgrass prairie and central mixed-grass prairie Bird Conservation Regions (BCR; Babcock et al. (1998)) cover portions of six sates in the central United States (NABCI 2020). Located in the rain-shadow of the Rocky Mountains, there is a strong west to east gradient in mean annual precipitation (average precipitation 29 - 102 cm/year) and a north to south gradient in mean temperature (average temperature 5 - 19 C) (Reese and Skagen 2017). Colloquially referred to as the “breadbasket of the America,” native prairie has been heavily converted to row crop agriculture due to the presence of the underlying High Plains aquifer (Comer et al. 2018; Haacker, Kendall, and Hyndman 2016; Steward et al. 2013).

The region hosts a diverse bird community, nearly 75% of these species have declining population trends making grassland birds the most imperiled bird community on the continent (Rosenberg et al. 2019). Grasshopper sparrows (*Ammodramus savannarum*) are a widely distributed prairie species, found through much of the United States east of the Rocky Mountains (Vickery 2020) and declining at 2.3% per year in the shortgrass ecoregion and 1.4% per year in the central mixed-grass ecoregion since 1966 (J. R. Sauer et al. 2020). In the shortgrass and central mixed-grass BCRs, grasshopper sparrows require large tracks (30 ha) of open grassland with areas of bareground and avoids woody encroachment (Vickery 2020).

### 2.2 Data

We used three different sources of data collected in 2017: IMBCR, BBS, and eBird (Figure 1). The IMBCR program uses a stratified hierarchical sampling design described in depth elsewhere (Pavlacky Jr et al. 2017), and here we summarize the details to provide sufficient context. Primary sampling units are 1 km^2^ grid cells selected at random from the sampling frame of each stratum, surveyed annually in the breeding season. Each primary sampling unit contains a 4 by 4 grid of 16 secondary sampling units (hereafter IMBCR points). These are point locations 250 m apart, placed more than 125 m from the primary sampling unit boundaries. On a survey, observers conduct six minute point counts at each IMBCR point in a primary sampling unit, recording distances to all detected birds or clusters of birds in one minute intervals. This protocol enables removal-in-time distance sampling, and estimation of density, detection, and availability. We expect that this is high quality data, with paid expert observers that are trained twice each year.

**Figure 1:**
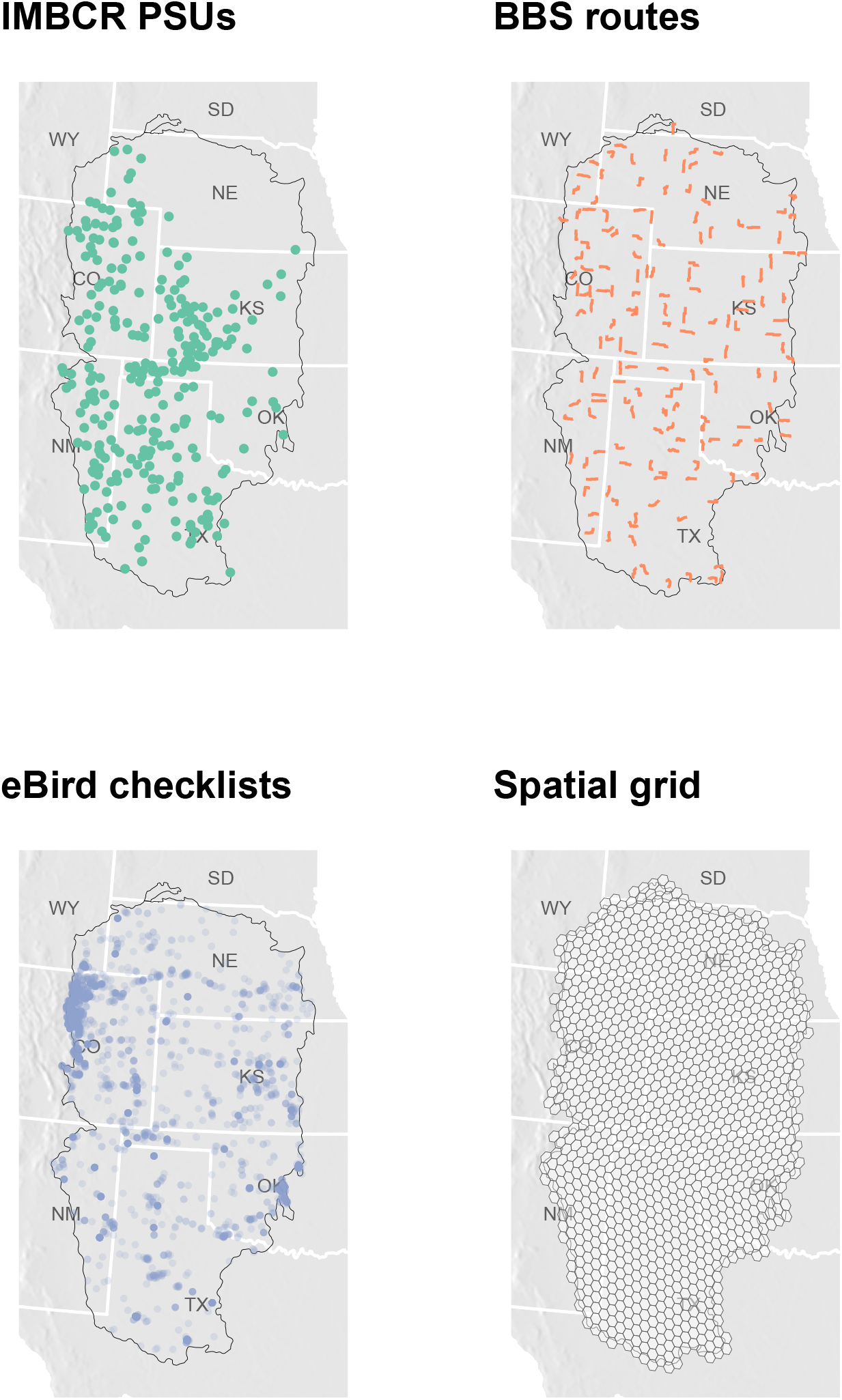
Location of IMBCR primary sampling units (PSUs) with 20 km spatial jitter to retain spatial anonymity, BBS routes, eBird checklists, and the hexagonal spatial grid used to model spatial autocorrelation.

The BBS data consist of counts of birds seen and heard within 400 m over a 3 minute time interval at 50 stops along each BBS route in the region. The BBS sampling frame is composed of 1 degree blocks of latitude and longitude (primary sampling units) that contain sufficient road length (Robbins and Van Velzen 1967).

The secondary sampling units are routes selected along roads within the primary blocks. Historically, blocks were selected using a random sampling design stratified by state, but now uses state by BCR stratification (John R. Sauer, Fallon, and Johnson 2003). Stops within routes are tertiary sampling units. A route consists of 50 roadside stops 800 m apart along a transect (a connected stretch of road) surveyed sequentially by an observer. Typically observers survey the same routes from year to year, and counts tend to be low for observers’ first year surveying a route (Kendall, Peterjohn, and Sauer 1996). While specific stop locations are not known precisely, route paths are available for 164 routes in the focal region. Thus we sum counts across stops within routes. The BBS uses experienced volunteers, and the resulting data are expected to be moderate or intermediate quality, suitable for regional conservation planning (Rosenberg et al. (2017); John R. Sauer et al. (2017), but see Thogmartin et al. (2006) and Sólymos et al. (2020)).

eBird data are community science observations generated by many different observers using various protocols. For consistency with IMBCR an BBS data we focus on stationary protocols only, where observers count all individuals encountered at a spatial location over a known time interval. These data contain a wide range of survey durations, and variable observer skill. Further, eBird data exhibit spatial bias in availability (Callaghan et al. 2019). We expect these data to be of mixed quality, less reliable than IMBCR or BBS data.

To explain local variation in density across the region, we extracted land cover data in a 125 m buffer from each IMBCR point from the National Land Cover Database. We summarized this data by computing the fraction of area within 125 m radius comprised of shrubland, grassland, and cultivated crop, generating one fractional summary per cover type for each IMBCR point. These three land cover summaries were used to explain local spatial variation in bird density for the IMBCR data.

Regional variation in density was modeled on a hexagonal grid over the spatial domain of the observed data, buffered by 50 km in every direction. The resulting grid has 1111 cells that are 779 km^2^ in area, spaced approximately 30 km apart. Spatial discretization is useful for constructing spatial random effects, because sparsity in the neighborhood structure permits computationally efficient representations of spatial autocorrelation (Besag and Kooperberg 1995). The size of the cells represents a tradeoff between computational efficiency and spatial grain. We selected cell sizes that led to reasonable computation times (< 1 day), and spatial scales similar to the largest sampling units (BBS routes, with all but one BBS route being completely contained in one cell and one BBS route spanning two cells).

### 2.3 Model structure

We consider multiple models that include auxiliary data with IMBCR distance sampling data. The IMBCR component is based on the distance sampling and removal-in-time model of Amundson, Royle, and Handel (2014), and we integrate auxiliary data using latent multivariate spatial fields, the “correlation approach” described in Pacifici et al. (2017). Models differ in terms of which auxiliary data augment IMBCR data.

#### 2.3.1 IMBCR model

We index secondary sampling points by *k*, so that we have point-transect data from points *k* = 1, …, *K*. When point *k* is surveyed, the radial distance and time-to-detection is recorded for each detected bird. The data consist of the total number of individuals detected at each point *y*_*k*_, along with the time period *t*_*i*_ and distance *d*_*i*_ for every detected individual *i* = 1, …, *n*, where *n* = ∑ _*k*_ *y*_*k*_. The original description of this model in Amundson, Royle, and Handel (2014) used an N-mixture model for abundance, and we use an equivalent integrated formulation (details in appendix):

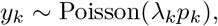

where *λ*_*k*_ is the expected population size in the sampled area around point *k*, and *p*_*k*_ is the probability that an individual is available and detected, given that it is present (Amundson, Royle, and Handel 2014). Expected population size is modeled as a function of known covariates and unknown spatial adjustments:

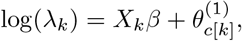

where *X*_*k*_ is a row vector of known covariates (a 1 to represent an intercept, and land cover fractions around point *k*), *β* is a vector of coefficients, and 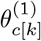 is a spatial adjustment corresponding to the hexagonal cell containing point *k*, denoted *c*[*k*]. Expected density can be computed by dividing by sampled area, e.g., expected number of individuals per square km at point *k* is would be 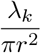 when using data in a radius of *r* km.

To be detected, individuals must be available (e.g., calling or visible), and perceived (e.g., heard or seen) by the observer. Represent the probability of availability as 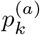 and the probability of perception conditional on availability be 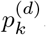. If availability is independent from perceptability, then 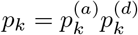 (Hostetter et al. 2019).

Individuals are only counted once during a survey, and the probability of availability at point *k* is the sum of the probabilities of availability in each of *T* time intervals:

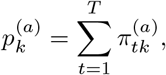

Where 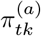 is the probability of availability in the *t*^*th*^ time interval (e.g., the *t*^*th*^ minute). These interval-wise probabilities are computed as:

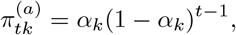

where *α*_*k*_ is the probability that an individual is available (e.g., calling and/or visible) during one time interval at point *k*. Heterogeneity in availability that can arise from survey conditions (e.g., temperature) can be incorporated in the model via *a*_*k*_ (Amundson, Royle, and Handel 2014). Here, we assume that *α* is constant, and remove the *k* subscript from subsequent notation.

Conditional on detection, the time interval observation model is:

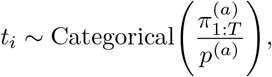

where 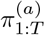 represents the vector containing elements 1, …, *T* (Amundson, Royle, and Handel 2014).

The probability of perception is modeled as a function of distance, discretised into 10 equidistant bins *b* = 1, …, *B* from 0 m to 125 m. The distance bin *d*_*i*_ of the *i*^*th*^ detected individual is modeled as:

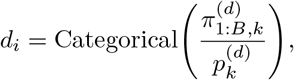

where 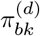 is the probability of being detected in distance bin *b* given availability, and 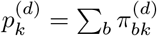. We use a half-normal detection function to obtain the cell probabilities (Amundson, Royle, and Handel 2014):

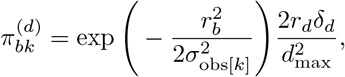

where *r*_*b*_ is the radial distance from the spatial coordinates of point *k* to the midpoint of distance class *b, σ*_obs_ is an observer-specific spatial decay parameter that represents how quickly the detection probabilities decrease as a function of distance (with *σ*_obs[*k*]_ denoting the particular value for the observer who surveyed point *k*), *δ*_*b*_ is the the width of distance bin *b*, and *d*_max_ is the maximum truncation distance for detections (Buckland et al. 2001).

By modeling the distance and time interval data separately we are assuming that they are (conditionally) independent. We assume no interaction between distance and time, such as would result if birds tended to move toward or away from observers once a survey begins. Following Amundson, Royle, and Handel (2014), we used ANOVA to test the assumption that distance and time of detection are independent. We failed to reject the assumption of independence (p = 0.28).

#### 2.3.2 BBS model

We use a simple count model for BBS data, aggregating counts among all 50 stops within each route. The BBS data consist of counts *z*_*r*_ at routes *r* = 1, …, *R*. These counts are modeled with a negative binomial distribution:

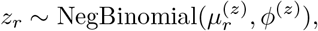

Where 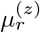 is the expected count at BBS route *r* and *φ*^(*z*)^ is an overdispersion parameter. The expected counts are a function of an overall mean and a spatial adjustment:

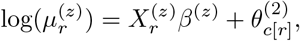

where 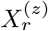 is a row vector of known covariates (a 1 to represent an intercept, and a binary indicator for whether it was an observer’s first time on a route (Kendall, Peterjohn, and Sauer 1996)), *β*^(*z*)^ is a parameter vector, and 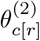 is a spatial adjustment.

#### 2.3.3 eBird model

The eBird model was similar in structure to the BBS model. The data consist of checklists *j* = 1, …, *J*, where each checklist has an associated count of the number of detected individuals, denoted *q*_*j*_. Again we use a negative binomial model:

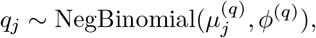

Where 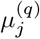 is the expected value, and *φ*^(*q*)^ is an overdispersion parameter. The expected value depends on covariates and varies spatially:

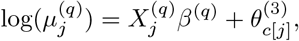

where 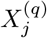 is a row vector of covariates (a 1 to represent an intercept, and a measure of “effort” or person-hours, which we compute as the product of duration and number of observers), *β*^(*q*)^ is a parameter vector, and 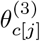 is a spatial adjustment.

#### 2.3.4 Sharing information among datasets

Correlations between the spatial adjustments allow for information to be shared among IMBCR, BBS, and eBird. Similar to Pacifici et al. (2017), we use a multivariate intrinsic conditional autoregressive prior for the spatial adjustments of cells *c* = 1, …, *C*. This allows for spatial autocorrelation and correlation among the three components simultaneously:

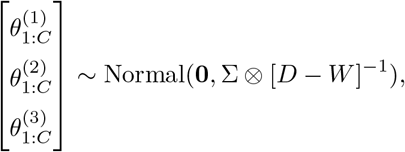

where **0** is a vector of zeros of length 3*C*, Σ is a 3 × 3 covariance matrix for IMBCR, BBS, and eBird data, ⊗ is the Kronecker product operator, *D* is a *C* × *C* diagonal matrix where *d*_*ii*_ is the number of neighbors of cell *i*, and *W* is a binary spatial neighbor adjacency matrix (Best, Richardson, and Thomson 2005).

### 2.4 Model comparison and evaluation

We randomly partitioned the set of 277 IMBCR primary sampling units (each of which contains up to 16 points) into a training, validation, and test set, using 142 of the transects for training, 67 for validation, and 68 for a test set. The training set was used for parameter estimation, the validation set was used for model comparison, and the test set was used for a final evaluation of generalization performance. We developed four models, and compared them all in terms of holdout log likelihood for the validation set:

- Model 1: IMBCR data only
- Model 2: IMBCR and BBS data
- Model 3: IMBCR and eBird data
- Model 4: (full model) IMBCR, BBS, and eBird data

Finally, to visualize model predictions we extracted covariate data on a spatial grid of points, representing cell centroids for a 1km^2^ cell area spatial grid over the study region.

### 2.5 Implementation

We implemented all models using dynamic Hamiltonian Monte Carlo in Stan (Carpenter et al. 2017). We used four chains, 4000 iterations per chain, and a maximum treedepth of 13. No divergent transitions or maximum treedepth warnings were encountered with these settings. Convergence was assessed using visual inspection of traceplots, and the R hat statistic. The Stan program corresponding to the full model is provided in the appendix.

## 3 Results

Both BBS and eBird data improved model performance on withheld IMBCR data, and the full model that used IMBCR, BBS, and eBird data together performed best on the validation set (Table 1). eBird and BBS data provided additional spatial information, increasing the posterior precision of the spatial random effects relative to the model trained only with IMBCR data (Figure 2). Spatial adjustments associated with eBird and BBS were weakly positively correlated with the spatial adjustments for the IMBCR submodel. The correlation between IMBCR and BBS adjustments was 0.19 (posterior median), with a 95% credible interval (CI) of (−0.26, 0.57). The posterior probability of a positive correlation between IMBCR and BBS adjustments was ≈ 0.79. The correlation between IMBCR and ebird adjustments was 0.29 (95% CI: −0.21, 0.7). The posterior probability of a positive correlation between IMBCR and eBird adjustments was ≈ 0.88. In contrast, BBS and eBird adjustments had a correlation of 0.45 (95% CI: 0.13, 0.71). The posterior probability of a positive correlation between BBS and eBird adjustments was ≈ 1. Indeed, the correlation between BBS and eBird spatial adjustments is probably higher than the correlation between IMBCR and BBS (posterior probability: ≈ 0.86), and between IMBCR and eBird (posterior probability: ≈ 0.7).

**Table 1:** Model performance on withheld validation data, sorted from best to worst.

| Model | Holdout log-likelihood |
| --- | --- |
| IMBCR + BBS + eBird | -1293 |
| IMBCR + BBS | -1300 |
| IMBCR + eBird | -1360 |
| IMBCR | -1371 |

**Figure 2:**
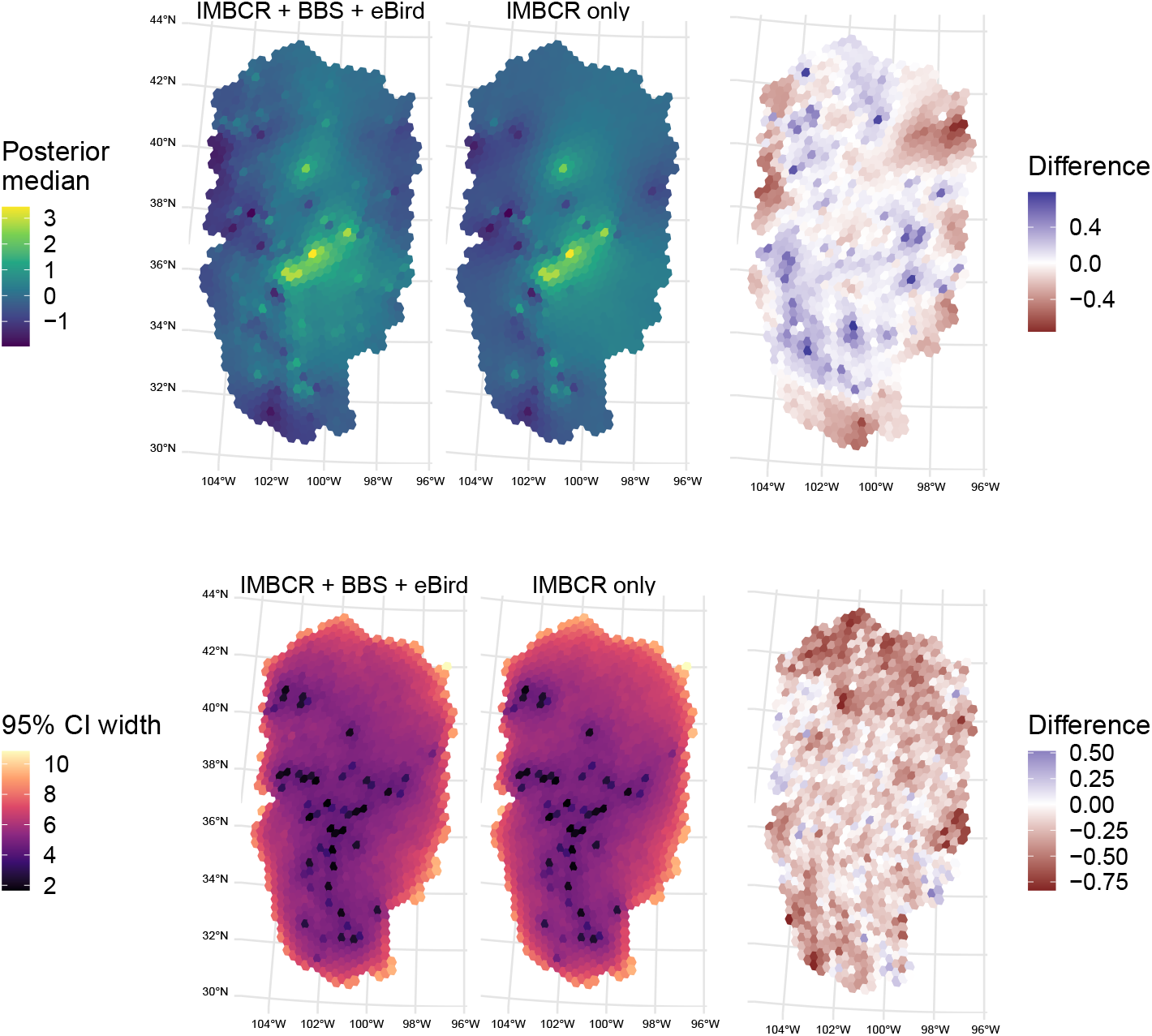
Comparison of spatial adjustments for and credible interval (CI) width for the full model and the model trained on IMBCR data only. The top row of plots display posterior medians for *θ*^(1)^, the spatial adjustments on IMBCR data from the full model (left) and the model using IMBCR data only (center), with the posterior median difference (right). Here, red indicates that the median for the full model was less than the median for the IMBCR only model. The bottom panel shows CI widths for the full and IMBCR only model, with differences shown on the right. Here, red indicates that the CI for the full model was narrower than the CI for the IMBCR only model. Values are shown on the natural log scale.

Grasshopper Sparrows were more abundant at points with a high fraction of grasslands, with a posterior median effect of 1.79 (95% CI: 0.94, 2.64). We did not observe as strong relationships between abundance and cultivated crop cover (posterior median: 0, 95% CI: (−0.8, 0.83)) or shrub cover (posterior median: 1, 95% CI: (−0.03, 1.8)) (Figure 3). As a consequence, the predicted densities and occupancy probabilities across the study region reflect both the large-scale spatial effects and fine-scale grassland cover effects (Figure 4).

**Figure 3:**
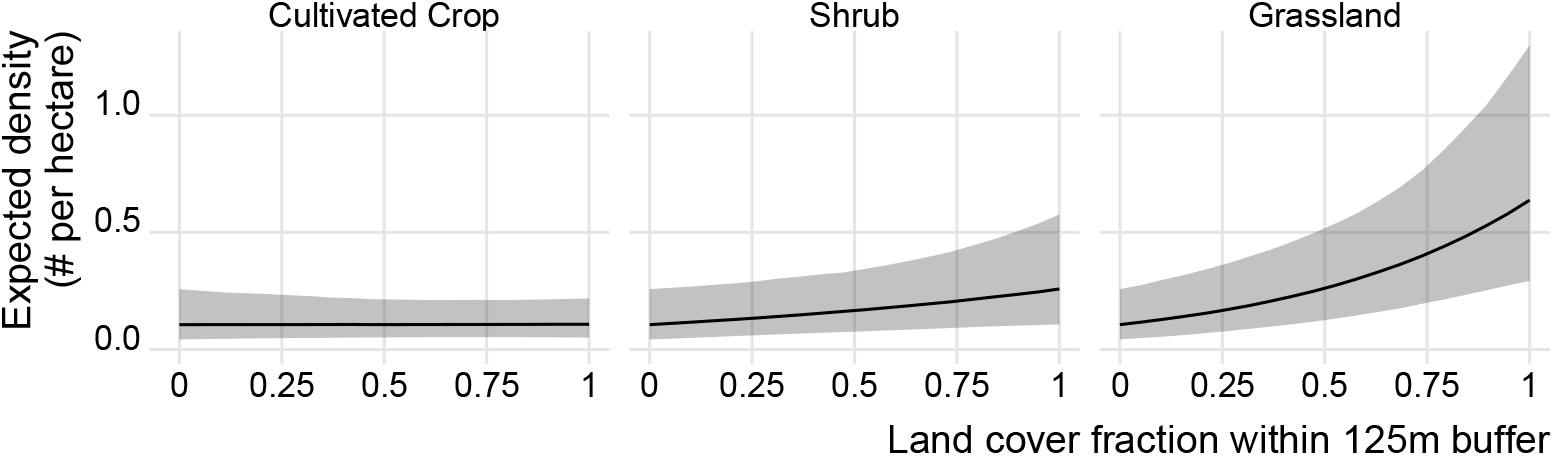
Expected density (individuals per hectare) as a function of fractional land cover within a 125 m buffer of an IMBCR point transect. Marginal effects are shown, fixing other land cover types to zero.

**Figure 4:**
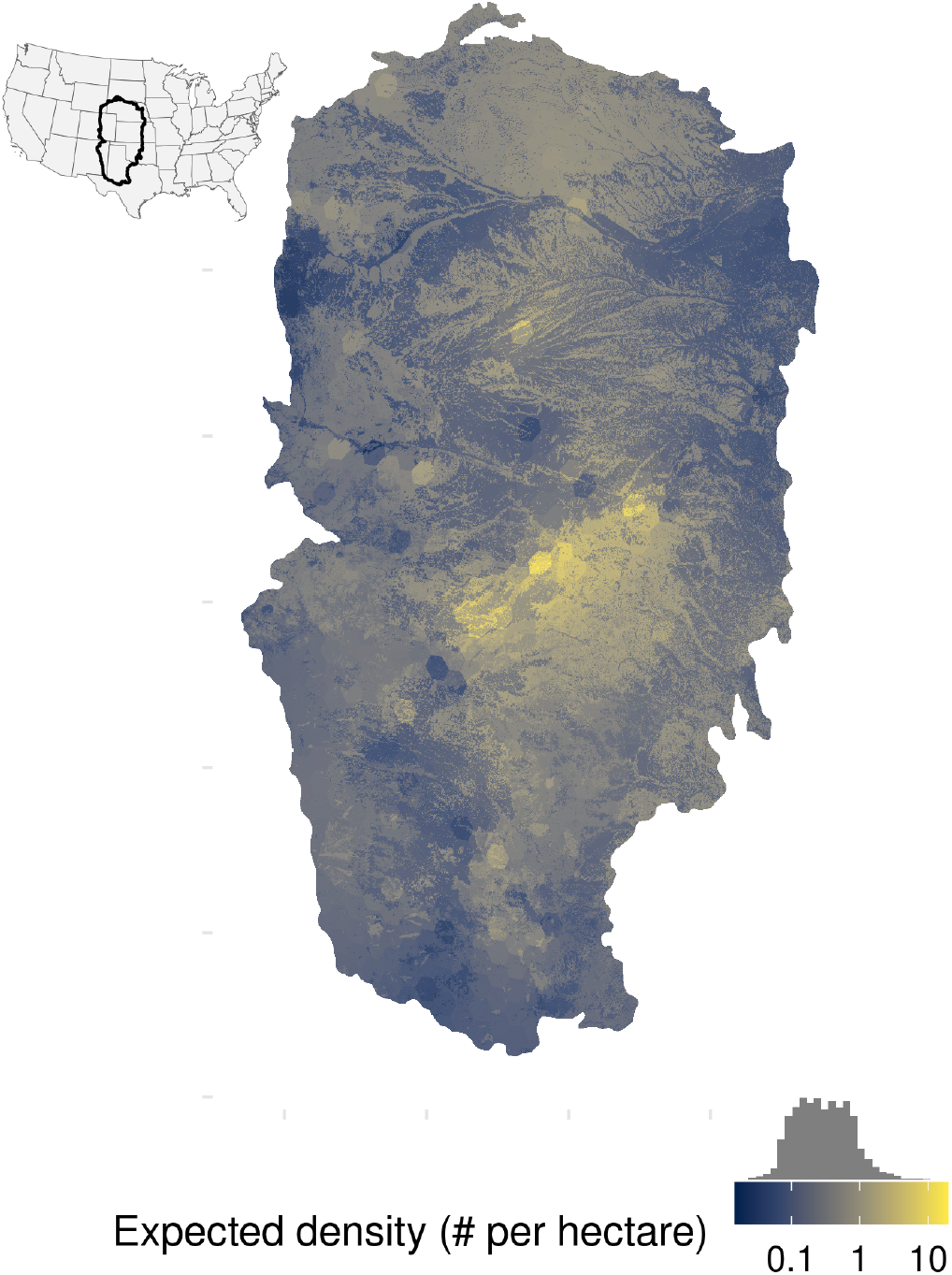
Spatial predictions of posterior median expected density across the study region.

Predictive performance of the final model on the test set was satisfactory. Empirical 95% credible interval coverage for counts in the test set was 99.7%. The fraction of zero counts in the test set was 0.60, and the 95% CI from the full model for the fraction of zeros in the test set was (0.48, 0.84) with a median value of 0.69. Similarly, the maximum count in the test set was 5, and the 95% CI for the maximum was (4, 80), with a median of 11.

Detection probabilities in the IMBCR point transect data varied by observers, such that the best observers were able to still detect calls at 125 m, and the worst observers had very low detection beyond 80 m (Figure 5). While it is not possible to separate abundance and detection effects in the BBS and eBird data, the covariates that we included did explain variation in expected counts. Specifically, we found a weak negative effect of a surveyor’s first year on a route in the BBS data. The posterior median for the first year effect on BBS counts was −0.35, and the 95% CI was (−1.1, 0.48). The posterior probability that this effect was negative was ≈ 0.82. We also found weak evidence for eBird counts being positively related to effort (posterior probability that the effect was positive: 0.8). The posterior median for the eBird effort effect was 0.1, and the 95% CI was (−0.13, 0.32).

**Figure 5:**
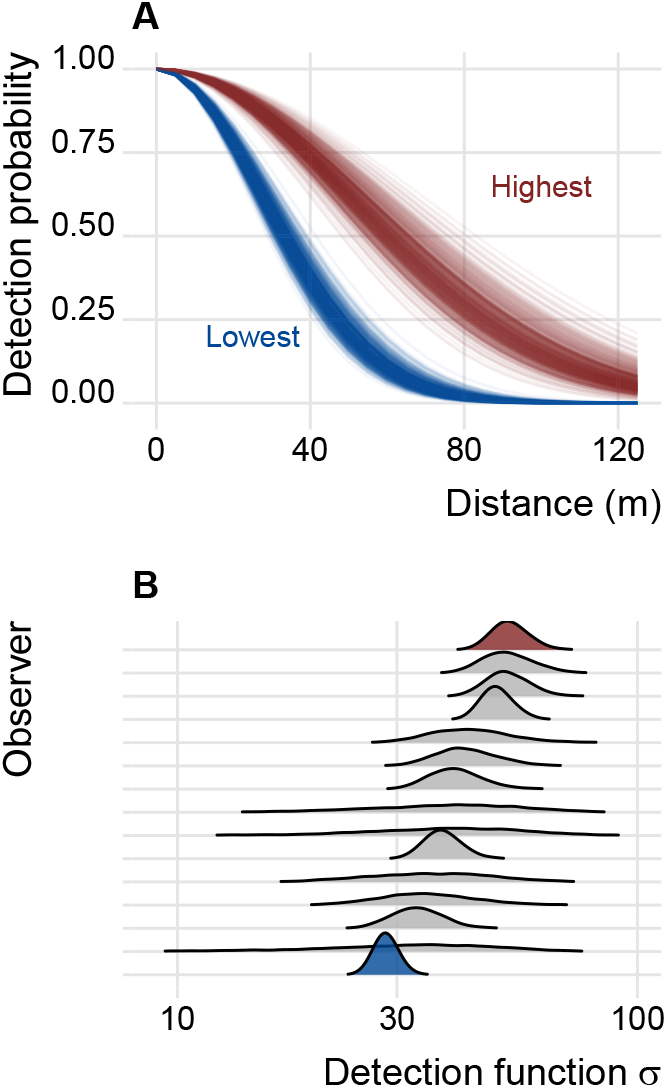
Differences in detection among observers. Panel A shows the estimated detection probability functions for the best (red) and worst (blue) observer at distances from 0 to 125 m, with one line per posterior draw. The y-axis in panel A represents the probability of detection given a call at a distance along the x-axis. Panel B shows the corresponding detection function standard deviation (*σ*) posteriors, with red and blue corresponding to the red and blue in panel A. Observers are sorted along the y-axis by the posterior means for *σ*.

## 4 Discussion

Here we show that both BBS and eBird data can provide useful information on bird abundance estimated using IMBCR distance sampling data, extending previous work that combined BBS and eBird data to estimate occupancy (Pacifici et al. 2017). Auxiliary data provided additional spatial information and increased the precision of spatial effects. Further, we showed that BBS and eBird are be more correlated with each other than they are correlated with IMBCR data. Despite this and despite considerable differences in sampling designs and survey protocols, we conclude that improved predictions of bird abundance can be obtained through a combination of data sources (Sólymos et al. 2020).

For some applications, abundance may be of interest in addition to occupancy, but data fusion for abundance estimation remains a developing research area. Recently Robinson et al. (2020) demonstrated predictive gains for observed counts when augmenting systematic survey data with eBird using random forest count models. A key difference between the model described in this paper and the approach of Robinson et al. (2020), is here we estimate density using a submodel that capitalizes on a removal-in-time distance sampling protocol that accounts for imperfect detection. However, machine learning and structured observation models are not mutually exclusive, and it would be possible to integrate the modeling approach described here with function approximators from machine learning Joseph (2020). Finally, we note that while we have focused on integrating different count datasets, there is a quickly expanding set of related methods being developed to use other data sources (such as capture-recapture data) with integrated population models (Zipkin and Saunders 2018).

To scale this fused abundance approach to a multi-species setting, there are some key future directions that stand out. First, to accommodate multiple species, it would be possible to extend the multivariate spatial field to include additional dimensions, e.g., three dimensions for each species, corresponding to IMBCR, BBS, and eBird spatial adjustments. However, this approach would quickly result in very high dimensional models as the number of species grows large. An alternative might use a convolution of two correlated spatial fields: one multivariate field with a dimension for each species, and another multivariate spatial field with a dimension for each data set (Majumdar and Gelfand 2007; Sollmann et al. 2016). Latent factor models could also reduce dimensionality (Hogan and Tchernis 2004; Ovaskainen et al. 2016). Hierarchical spatial embeddings might provide an even more scalable solution, and have already proved useful for massive multi-species models of BBS data (Joseph 2020). Finally, while we have used a discrete areal spatial representation, treating space continuously may be another valid option, particularly given recent progress on approximate Gaussian process models for larger spatial data (Tikhonov et al. 2020).

Other extensions could reveal temporal dynamics. Dynamic occupancy models stand out as an immediate choice for estimating colonization and extinction probabilities in space (Royle and Kéry 2007). Temporal abundance models could leverage process-based dynamical models (Dail and Madsen 2011; Hostetler and Chandler 2015), statistical autocorrelation (Wilson, Smith, and Naujokaitis-Lewis 2018), nonlinear dynamics (Clark and Luis 2020), and/or spatiotemporal information (Chevalier and Knape 2020).

We chose to integrate data using correlations rather than a joint likelihood because of differences in survey protocols and spatial locations (Miller et al. 2019). The correlation approach can limit bias that may arise from aggregating non-standardized data sources (Pacifici et al. 2017). Because the BBS is a roadside survey and the specific stop locations along each path is unknown, it was not clear how much information would be contained in the BBS data to inform IMBCR counts. If we had the exact spatial locations of each of the 50 stops, we would have felt more comfortable modeling counts at the stop level, as the spatial scale of stops is similar (same order of magnitude detection radius) to that of the IMBCR data. Similarly, for eBird data, it was not clear whether a joint likelihood would help, given the mixed quality of these community science observations, and the fact that the maximum radius at which birds would be detected and recorded is unknown and could vary among checklists (Pacifici et al. 2017). Finally, we opted not to use a covariate approach, because our eventual goal is to build a model to predict abundance forward in time, and in this setting, all of these data would be unknown such that neither the BBS or eBird data could be treated as observed covariates.

While the current approach borrows spatial information from BBS and eBird to inform the IMBCR submodel, it does so using a single correlation parameter that is applied across the entire region. This might not be ideal, given that the BBS is a roadside survey, and the eBird checklists tend to concentrate in areas of high population density and/or around major interstate roads (Figure 1). This correspondence might underlie the relatively strong positive correlations between BBS and eBird reported here. As a consequence, it may be that these auxiliary data are most informative for IMBCR data from similar locations (e.g., point transects near roads). Future work might evaluate whether there are advantages to allowing for more flexible spatial correlation structures, that might share more or less information among datasets depending on spatial similarity. A “covariate” or conditional data integration approach with spatially varying coefficients might be particularly amenable to this (Hanks, Hooten, and Baker 2011; Pacifici et al. 2017), to allow the relationship between auxiliary and focal data to depend on land cover and road density (Finley 2011).

A variety of data are available to understand the distribution and abundance of birds, but integrating different datasets remains a key area for development (La Sorte et al. 2018). Here, we demonstrate that the combination of IMBCR, BBS, and eBird data provides better predictive performance than using IMBCR data alone. Having outlined some future directions for multi-species and multi-temporal settings, we hope that future work can integrate additional data to improve estimates of bird abundance, understand ecological processes, and improve conservation outcomes.

## 5 Acknowledgements

We would like to thank John Sauer for discussions about the BBS model, and Chris Latimer for helping with processing BBS route data. We would also like to thank Jennifer Balch for help with soliciting funding for this work. This work was supported in part by the University of Colorado Boulder’s Grand Challenge initiative and their investment in Earth Lab.

## 7 Appendix

The point transect model described here is equivalent to the original model of Amundson, Royle, and Handel (2014), which has a hierarchical three stage model for the number of detected animals:

- First stage: latent true number of individuals *N*_*k*_ ∼ Poisson(*λ*_*k*_), where *λ*_*k*_ is the expected number of individuals at point *k*.
- Second stage: latent number of available individuals *a*_*k*_ ∼ Binomial(*N*_*k*_, *p*^(*a*)^), where *a*_*k*_ is the number of individuals available for detection at point *k* and *p*^(*a*)^ is the probability of availability.
- Third stage: number of detected individuals *y*_*k*_ ∼ Binomial(*a*_*k*_, *p*^(*d*)^), where *p*^(*d*)^ is the probability of detection conditional on availability.

We can collapse these three stages into one. First, recognize that this is a hierarchical Poisson-binomial-binomial compound distribution The last two binomial components can be collapsed into one using the binomial-binomial hierarchy, to obtain a marginal distribution for *y*_*k*_:

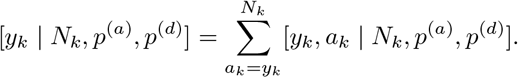

This factorizes:

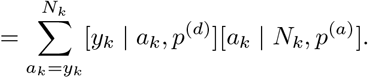

Plugging in the binomial probability mass functions:

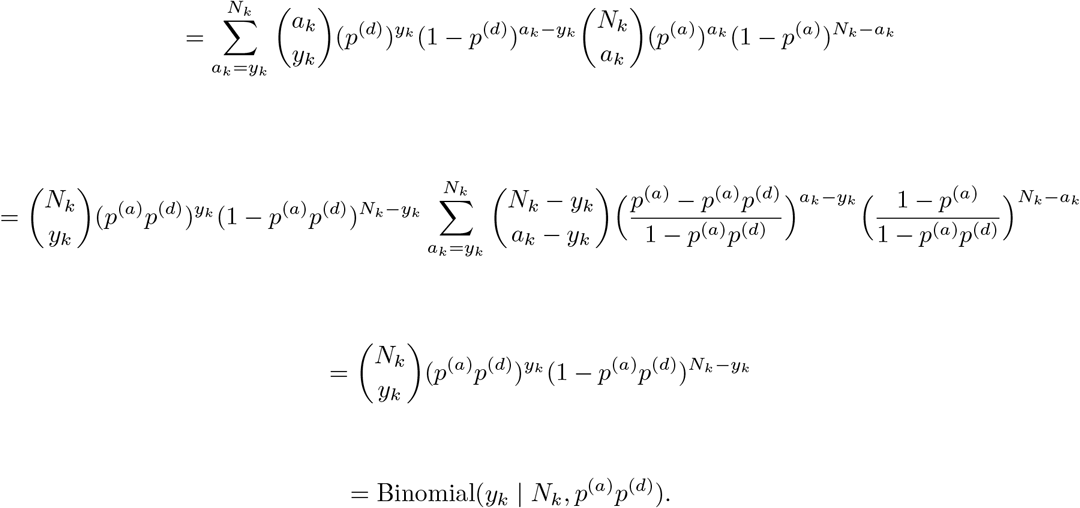

Now that we have collapsed the problem from a Poisson-binomial-binomial hierarchy to a Poisson-binomial hierarchy, we can further simplify to obtain the marginal distribution of the observed number of individuals at point *k*. If *N*_*k*_ ∼ Poisson(*λ*_*k*_) and *y*_*k*_ ∼ Binomial(*N*_*k*_, *p*^(*a*)^*p*^(*d*)^), the Poisson-binomial hierarchy implies that the marginal distribution of *y*_*k*_ is Poisson (Casella and Berger 2002):

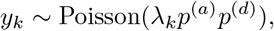

which is denoted *y*_*k*_ ∼ Poisson(*λ*_*k*_*p*) in the methods section, collapsing the two detection terms into one (*p* = *p*^(*a*)^*p*^(*d*)^). This marginal distribution is useful, because *y*_*k*_ is observed, and writing the model in this form removes all discrete latent parameters. Thus, the log density of the model is differentiable, enabling modern Hamiltonian Monte Carlo sampling from the posterior. We implemented all models in the Stan probabilistic programming language (Carpenter et al. 2017), and provide the full model code for reference below:

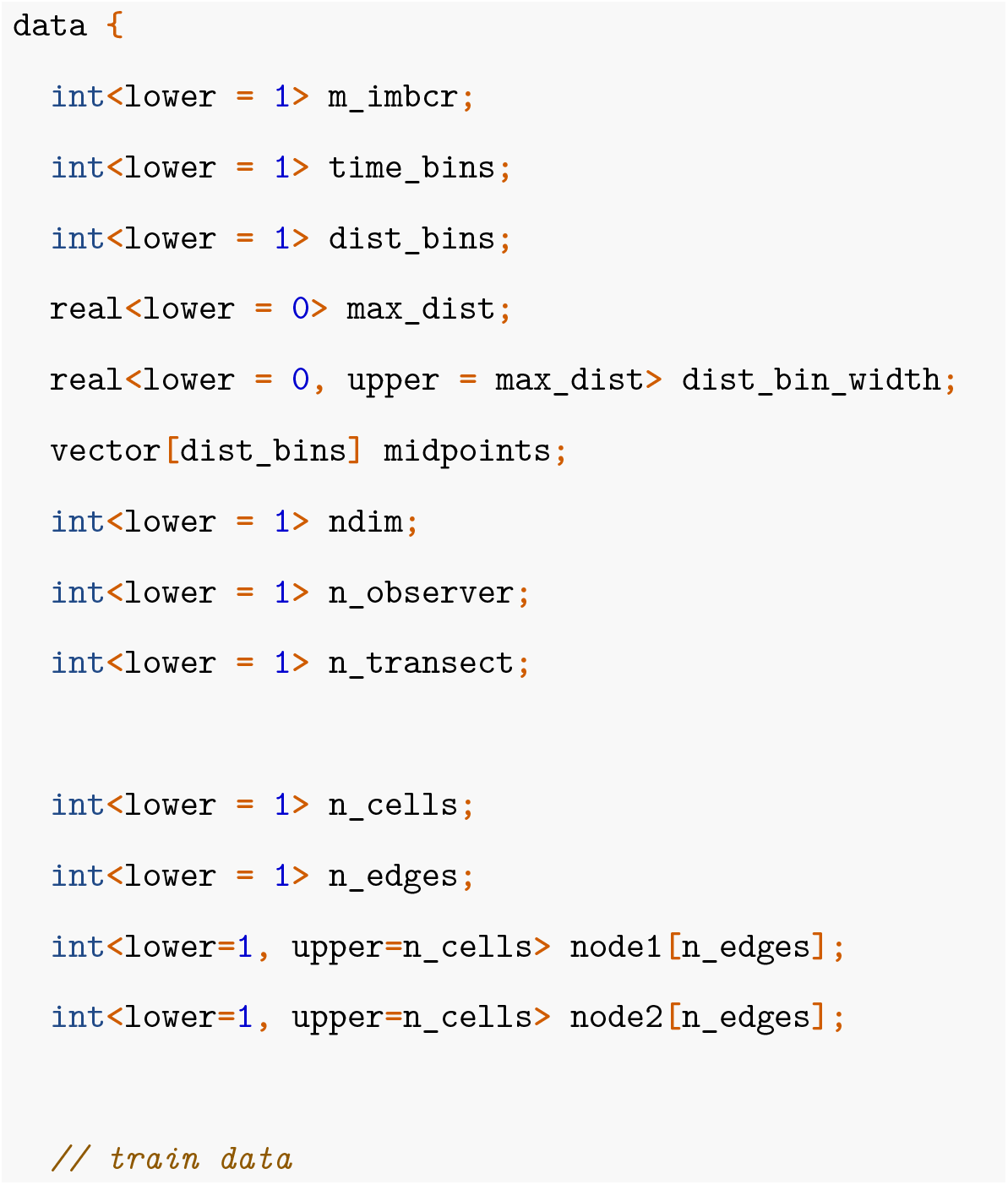
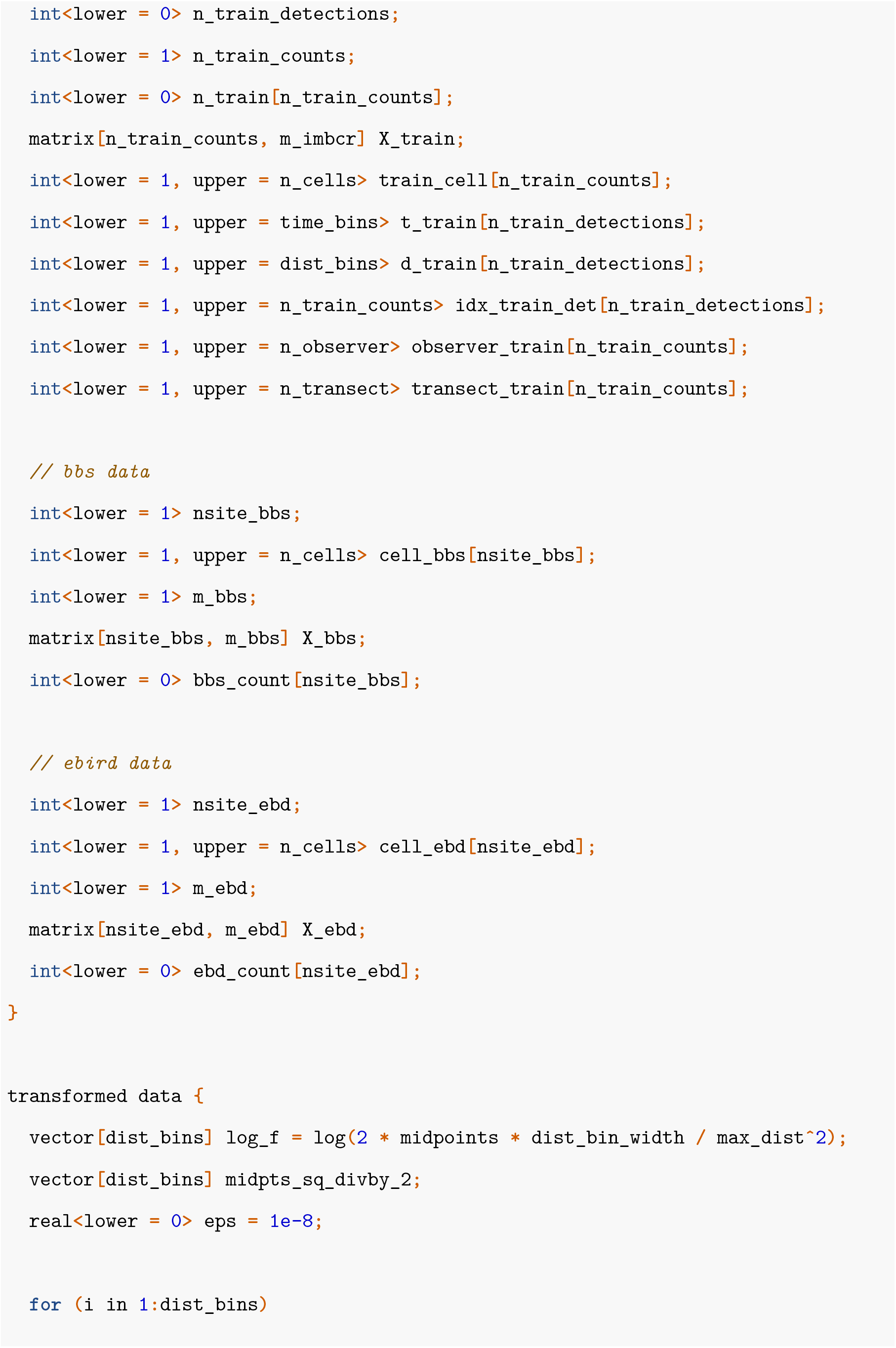
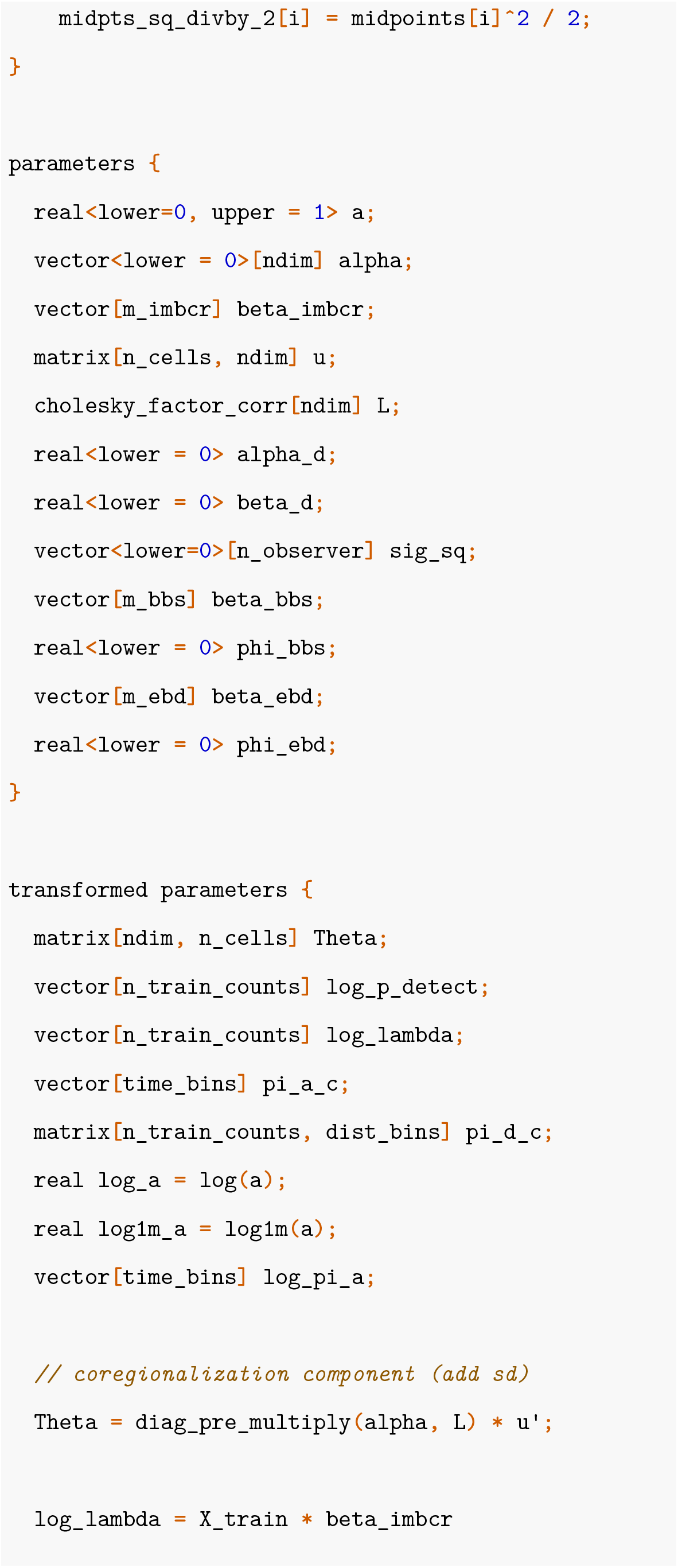
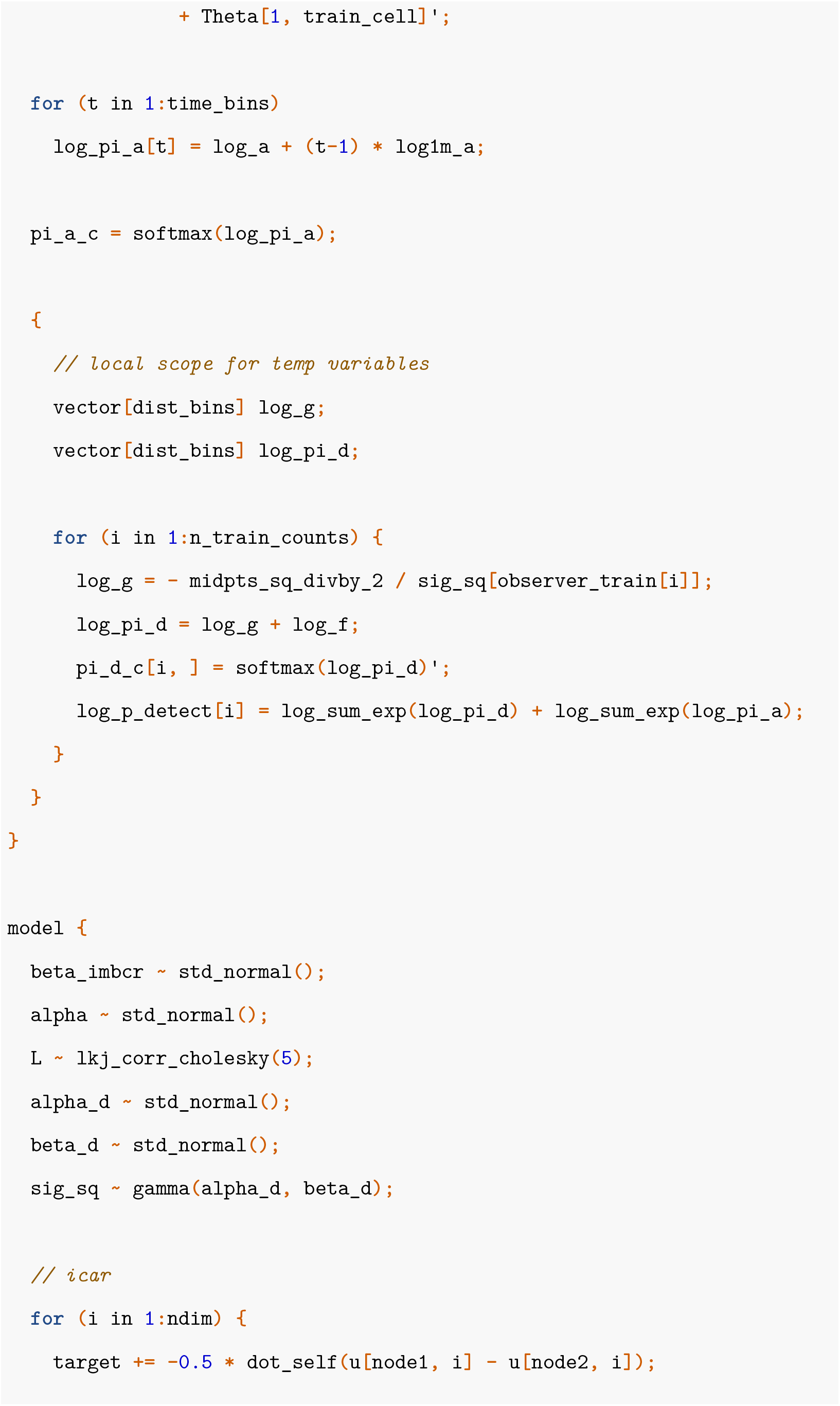
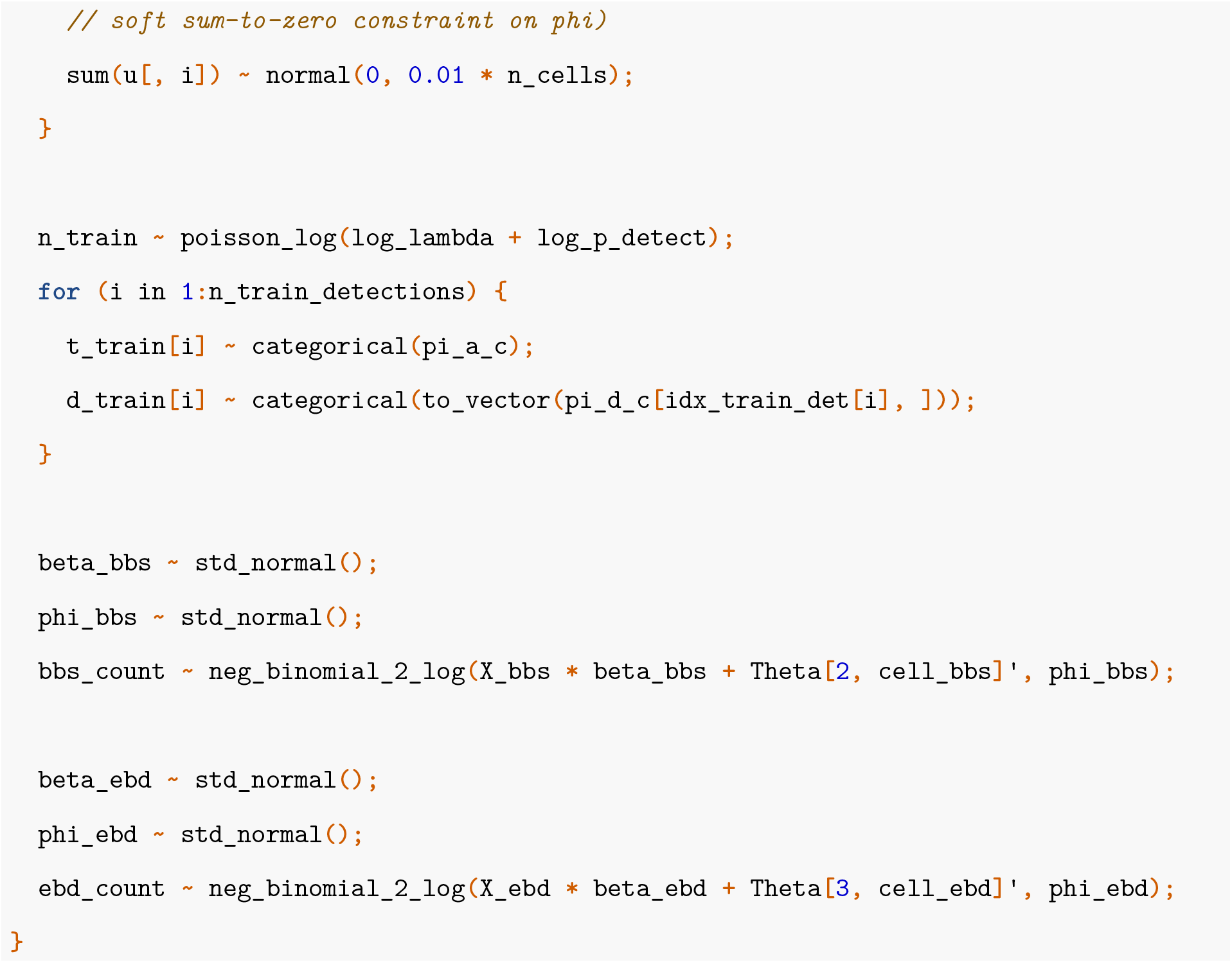

